# Sensory-motor brain dynamics reflect architectural affordances

**DOI:** 10.1101/520080

**Authors:** Zakaria Djebbara, Lars Brorson Fich, Laura Petrini, Klaus Gramann

## Abstract

Anticipating meaningful actions in the environment is an essential function of the brain. Such predictive mechanisms originate from the motor system and allow for inferring actions from environmental affordances, the potential to act within a specific environment. Using architecture, we provide a unique perspective to the abiding debate in cognitive neuroscience and philosophy on whether cognition depends on movement or is decoupled from our physical structure. To investigate cognitive processes associated with architectural affordances, we used a Mobile Brain/Body Imaging approach recording brain activity synchronized to head-mounted virtual reality. Participants perceived and acted upon virtual transitions ranging from non-passable to easily passable. We demonstrate that early sensory brain activity, upon revealing the environment and before actual movement, differed as a function of affordances. Additionally, movement through transitions was preceded by a motor-related negative component also depended on affordances. Our results suggest that potential actions afforded by an environment influence perception.

**SIGNIFICANCE STATEMENT:** By using electroencephalography and virtual reality, our research provide a unique perspective to the centurial open-ended debate in cognitive neuroscience and philosophy on the relation between cognition, movement and environment. Our results indicate that cortical potentials vary as a function of bodily affordances reflected by the physical environment. Firstly, the results of this study implies that cognition is inherently related to potential movement of the body, thus we advance that action is interrelated with perception, actively influencing the perceivable environment. Secondly, as cortical potentials are influenced by the potential to move, which in turn is the task of architectural design, architects holds largely a privilege of human health, and thus potentially capable of provoking and preventing physiological conditions.

## INTRODUCTION

The affordance of a given spatial environment, defined as the perception of possibilities for, or restraints on, action that the environment offers, is essential for an agent to produce meaningful behavior. Thus, the affordances of the spatial environment becomes a central concept for humans interacting with their world. The term *affordances* was first introduced by Gibson (1), and later specified by various authors including Clark who defines affordance as “[…] *the possibilitiesfor use, interventionandactionwhich the physical world offers a given agent and are determined by the ‘fit’ between the agent’s physical structure, capacities and skills and the action-related properties of the environment itself.”* (2). In light of emerging theories of embodied cognition, the perception of the environment may be dependent on proprioceptive mechanisms. According to predictive processing, a neuroscientifically based theory of embodied cognition (3–5), motor systems, similar to perceptual processes, aim at cancelling out continuously incoming bottom-up sensory signals with top-down predictions. In this perspective, movement emerges as a result of an *active inference* that attempts to either minimizing motor trajectory prediction errors by acting, and thus perceiving the unfolding of the predicted movement, or by changing perception itself (6–8). From the standpoint of active inference, motor systems suppress errors through a dynamic interchange of prediction and action. In other words, there are two ways to minimizing prediction errors; one is to adjust predictions to fit the current sensory input, while another is to adapt the unfolding of movement to make predictions come true. It is a unifying perspective on perception and action suggesting that action is both perceived and caused by perception (9). Hence, action, perception, and cognition coordinate to move the body in ways that conform a transitional set of expectations (10). The claim we seek to investigate in the present study is that perception is rooted in action, creating an action-perception loop, informed by dynamically (top-down/bottom-up) generated prediction errors. Ultimately, the argument is that perception is not the sole result of sensing the physical world, but unfolds as an ongoing interaction between sensory processes and bodily actions. Such a claim has philosophical and neuroscientific significance as the neural dynamics underlying perception would be intimately dependent on the affordances of a given environment.

To further investigate this claim, we used electroencephalographic (EEG) recordings to address the neural dynamics of action-perception interactions through affordance manipulations in architectural experiences. More specifically, we investigated the affordances of transitions as they form an ideal candidate due to their dynamic nature concerning the duration of altering one condition to another (11). We here confine transitions to *the passage between spaces* which, according to the enactivists’ proposed action-perception loop, will be experience dependent on the affordances offered by the passage itself. From an architecturally historical point of view, the use of transitions have evidently been exploited at least since eleventh-thirteenth dynasties (e.g., Fazio et al., 2008, chaps. 1, 2, 5). Written interest in human experience of architectural settings has been established at least for the last two millennia (e.g., Norberg-Schulz, 1965; Palladio, 1997; Pallasmaa, 2011; Rasmussen, 1959; Vitruvius and Morgan, 1960). Despite transitions being ubiquitous in architecture, the underlying mechanisms of how transitions affect human perceivers appears to have taken an implicit, overlooked, and close to nonexistent position in architectural discourse, with few exceptions (15, 18–20). Due to the dynamic nature of architecture, an essential part of transitions and experiencing architecture is that of being able to act (21). Traditionally, investigations of architectural experiences are phenomenological - the description of phenomena in how experience gives access to a world of space and time (14, 22–24). Such descriptions find specifically movement of the individual to be an expression of a holistic experience of architecture (14, 22), linking the nature of movement to architectural experiences (25). Transitions in architecture depend on voluntary movement and thus a prerequisite for any transit is a goal, which in turn calls for action planning. Coarsely three parameters compose a transition: a motivated goal, a change in physical environment and the unfolding of action. All three parameters are interdependent, as reaching a goal depends on the affordance offered by an environment, and also propels the body in space contributing to experience. Architectural transitions thus include the attenuation of an agent’s experience through movements and how such movements animate the body through environmental changes.

Data from neuroscientific experiments addressing this issue might contribute to discussions centered on philosophical questions on how we relate to the world. For long, enactivists have implicated the reciprocal dependency of the living organism, as a self-organized living system, and the embedded body in a world for cognition (26–28). Enactivism is rooted in phenomenology (21, 29), similar to prominent architectural theorists, who put body, action, and cognition central to experience. Active inference closely relates to enactivism, in the sense that we act to perceive, and vice versa. Such a thesis rests on a hierarchical and dynamic model of the world, which temporally dissociates lower sensorimotor inferences from higher motivated goals, as fast and slow, respectively (30). Fast, lower sensorimotor inferences depict processes of affordances, which thereby must be present in early stages of perception. Hierarchical affordance competition (HAC; Pezzulo and Cisek, 2016) takes the temporal aspect of affordances much further, by suggesting that cortical activity relates to the immediate decision of action selection, which occurs fluently during movement. Such an account of temporally extended affordance is in accordance with active inferences.

To investigate the impact of environmental affordances on early sensory processing in actively transiting humans, we used a Mobile Brain/Body Imaging approach (32–34) recording brain activity with EEG synchronized to movement recordings and head mounted virtual reality (VR). This approach allows for investigating brain dynamics of participants perceiving an environment and the transitions contained therein as well as brain dynamics during the transitions itself. Previous studies investigating event-related potential (ERP) activity in stationary participants demonstrated slow cortical potentials to indicate anticipative motor behavior (for an overview, see Luck and Kappenman, 2011, chap. 8). Known motor-related cortical components (MRCPs) are the readiness potential (RP; Kornhuber and Deecke, 2016), contingent negative variation (CNV), and the stimulus-preceding negativity (SPN; Brunia, 2003), which can be seen as indicators of predictive behavior (38). MRCPs are negative going waveforms preceding an actual, or imagined, motor execution. However, these negative components are associated with multiple processes including sensory, cognitive, and motor systems. In a study by Bozzacchi et al. (39), the authors attempted to measure affordances of a physical object by evaluating whether the anticipated consequence of action itself influence the brain activity preceding a self-paced action. The authors compared MRCPs of situations where it was possible to reach out and grasp a cup, versus situations where it was impossible to grasp the cup, by tying the hands of the participants. A motor execution was forced at all times. In situations where it was impossible to grasp the cup, the authors reported an absence of early activity over the parietal cortex, and found instead increased activity over the prefrontal cortex. The results were interpreted as reflecting an awareness of the inability to execute a goal-oriented action. Closely related to the MRCPs is the post-imperative negative variation (PINV),a negative going waveform that is present succeeding an imperative stimulus. It reflects the immediate motor execution related to the onset of an imperative stimulus and was observed during experiments investigating learned helplessness or loss of control (40, 41). The PINV thus allows linking of motor relatedpotentials to anticipation of affective states (42).

If an enactive account of perception, action and cognition is correct, affordances intimately relate to higher hierarchical levels through low-level perceptual cues. Such an account would situate processing of affordances at a similar stage as early perceptual processes and should reveal differences in sensory and motor-related ERPs associated with the perceived affordance of an environment. To investigate whether brain activity is altered depending on affordances offered by the environment, we presented human observers with environmental stimuli that allowed or prohibited a transition from one room to the next. To this end, participants were presented with a view into a room containing one door of different widths, allowing or prohibiting a transition into the next room and thus providing different affordances. We expected to find differences in cortical responses to co-vary as a function of affordances over sensory and motor areas. In addition, we expected differences in motor-related cortical potentials as a function of the environmental affordances when participants were instructed to walk through the door or to remain in the same room.

## METHODS

### Participants

20 participants (9 female) without history of neurological pathologies were recruited from a participant pool of the Technical University of Berlin, Berlin. All participants read and signed a written informed consent about the experimental protocol, which was approved by the local ethics committee. Participants received either monetary compensation (10€/hour) or accredited course hours. The mean age was 28.1 years (σ = 6.2), all participants had normal or corrected to normal vision and none had a specific background in architecture (no architects or architectural students). One participant was excluded due to technical issues of the experimental setup.

### Paradigm description

The experiment took place in the Berlin Mobile Brain/Body Imaging Laboratories (BeMoBIL) with one of the experimental rooms providing a space of 160 m^2^. The size of the virtual space was 9 × 5 meters with a room size of 4.5 × 5 meters for the first room and a room size of 4.5 × 5 meters for the second room. Participants performed a forewarned (S1-S2) *Go/NoGo* paradigm (pseudo-randomized 50/50) in the virtual reality environment that required them to walk from one room to a second room. Doors of different width ranging from unpassable (20 cm, *Narrow*) to passable (100 cm, *Mid*) to easily passible (1500 cm, *Wide*) manipulated the transition affordance between rooms. The experiment consisted of a 3 × 2 repeated measures design including the factors door width (*Narrow, Mid, Wide*; pseudo-randomized) and movement instruction (*Go*, *NoGo*). A total of 240 trials per participant was collected with 40 trials for each of the factor levels. One trial consisted of a participant starting in a dark environment on a predefined starting square (see Figure 1). The “lights” would go on after a random inter-trial-interval (mean = 3 s, σ = 1 s), and participants faced a room with a closed door. They were instructed to wait (mean = 6 s, σ = 1 s) for a color change of the door with a change to green indicating a *Go* trial and a change to red indicating a *NoGo* trial. In case of a green door, the participant walked towards the door, which would slide aside. Upon entering the subsequent space, participants were instructed to find and virtually touch a red rotating circle by using the controller. The circle would inform the participant to have earned another 0.1€ to their basic reimbursement of 10 Euro per hour. After each trial, participants had to give an emotional rating for the environment irrespective of whether they transitioned through the door (*Go* condition) or whether they remained in the same room (*NoGo* condition) without transition. To this end, participants were instructed to go back to the starting square, and fill in a virtual Self-Assessment Manikin (SAM) questionnaire, using a laser pointer from the controller, and to subsequently pulling the response button located at the pointer finger to turn the “lights off”. The lights would go back on automatically to start the next trial.

In *Go*-trials, participants were instructed to walk towards the door and into the second room even in case the door was too narrow to pass. This was done to control for motor execution in the *Go*-condition and to allow movement towards the goal irrespective of the affordance (passable vs. unpassable). Upon touching the surrounding walls, the walls would turn red and inform the participants they have failed to pass, and thus must return to the start square, fill in the virtual SAM and start the next trial by pulling the trigger. Participants would quickly notice that the narrow door (20 cm) was impossible to pass without producing the warning feedback that they have failed to pass. All participants had a training phase to get accustomed to the VR environment and the different conditions. The experimenter observed the participants from a control room, separated from the experimental space, using two cameras and a mirrored display of the virtual environment to reduce interactions to a minimum during the experiments.

**Fig. 1.**
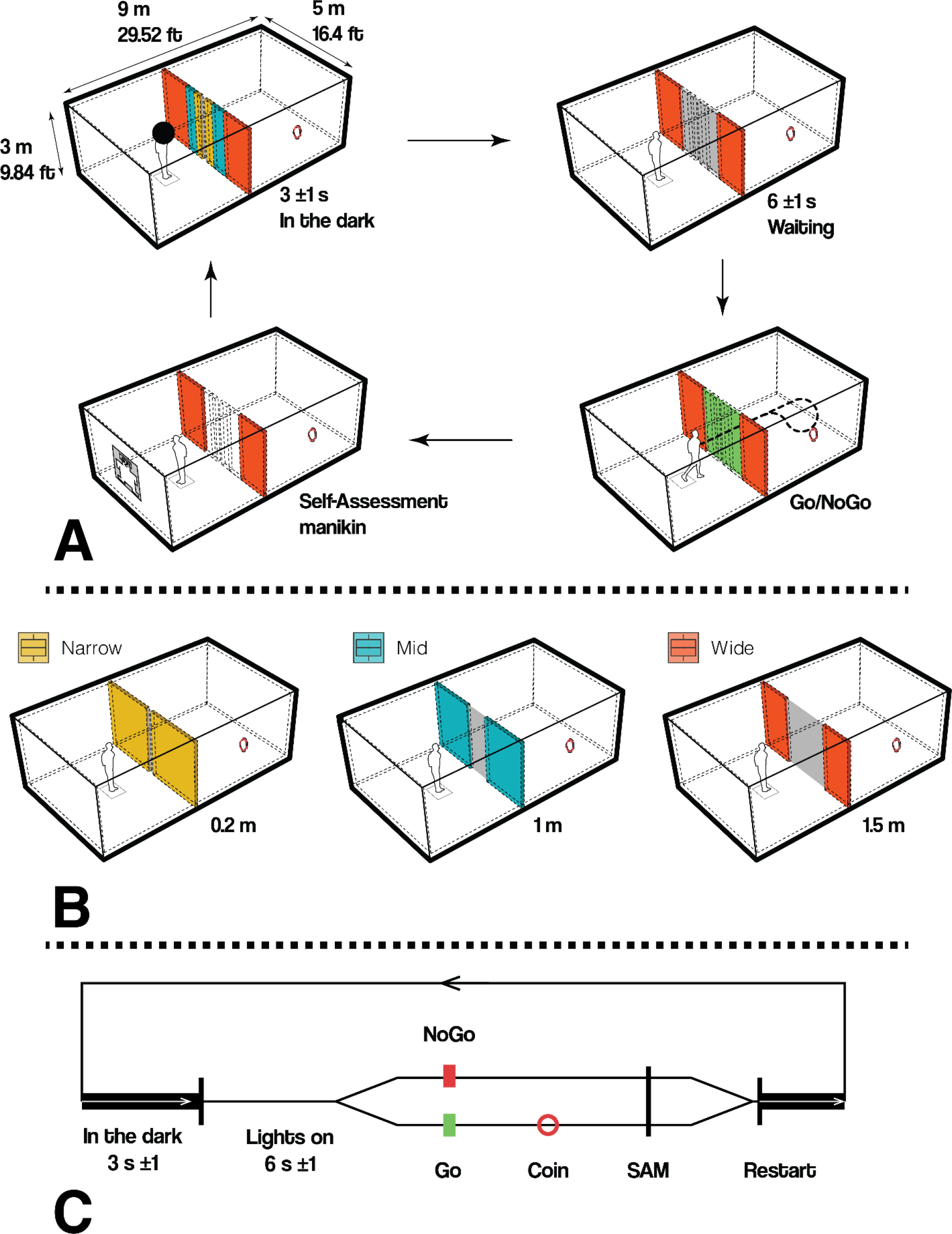
**(A)** Participants were instructed to stand in the start-square. A black sphere would restrict their vision to pure black for 3 seconds, σ = 1. The moment the black sphere disappears, participants perceive the door they have to pass. They wait for the imperative stimulus, either a green door (Go) or a red door (NoGo), for 6 seconds, σ = 1. In case of Go, participants were instructed to pass the opening, virtually touch the red circle, which in turn would release a monetary bonus, return to start square and answer the virtual SAM questionnaire. In case of NoGo, participants were instructed to turn around and answer the virtual SAM. **(B)** The three different doors were dimensioned as following *Narrow* 0.2 meter, *Mid* 1 meter and *Wide* 1.5 meters. Note the color code for each door as they are used throughout the paper. **(C)** The diagrammatic timeline depicts a the sequences of events for a single trial in conceptual manner.

### Subjective and Behavioral data

To investigate the subjective experience of the transitions, we introduced the participants with a virtual Self-Assessment Manikin (SAM) questionnaire after each trial. The SAM is a pictorial assessment of pleasure, arousal and dominance on a 5-point Likert scale (43). The manikin display ranges from smiling to frowning (*pleasure*), from a dot in the stomach to an explosion (*arousal*) and from being very small to very big (*dominance*). Participants were asked to self-assess their current state after each trial. Furthermore, we measured the reaction time from the onset of the *Go*-stimulus (door color change) to reaching the opening-threshold itself, to assess the behavior. The data was analyzed using ANOVA with the width of the doors as repeated measures factor. In case of violation of normality and homogeneity, corrected p-values are reported. For post-hoc analysis, the data was contrasted using Tukey HSD.

### EEG Recording and data analysis

To investigate the impact of transitional affordances on human cognition and brain dynamics, we used a MoBI approach (32–34, 44) recording human brain dynamics in participants actively transitioning through virtual rooms. All data streams were recorded and synchronized using LabStreamingLayer (LSL; Kothe, 2014). Participants wore a backpack, which held a high-performance gaming computer to render the VR environment (Zotac, PC Partner Limited, Hong Kong, China) attached to two batteries and an EEG amplifier system. We combined a Windows Mixed Reality (WMR; 2.89”, 2880 × 1440 resolution, update rate at 90 Hz, 100 degree field of view with a weight of 440 grams, linked to the Zotac computer through HDMI) headset and one controller by ACER to display and interact with the virtual environment based on Unity (see Figure 2). Events for recordings of performance and physiological data were triggered by the position of the participant in the tracking space or by the respective response buttons of the remote control. Specific events, such as touching the wall, all button presses, transitioning through the door, answering the questionnaire and all cases of “lights on” (and off), were synchronized with the recorded brain activity and the presented VR environment through LSL.

**Fig. 2.**
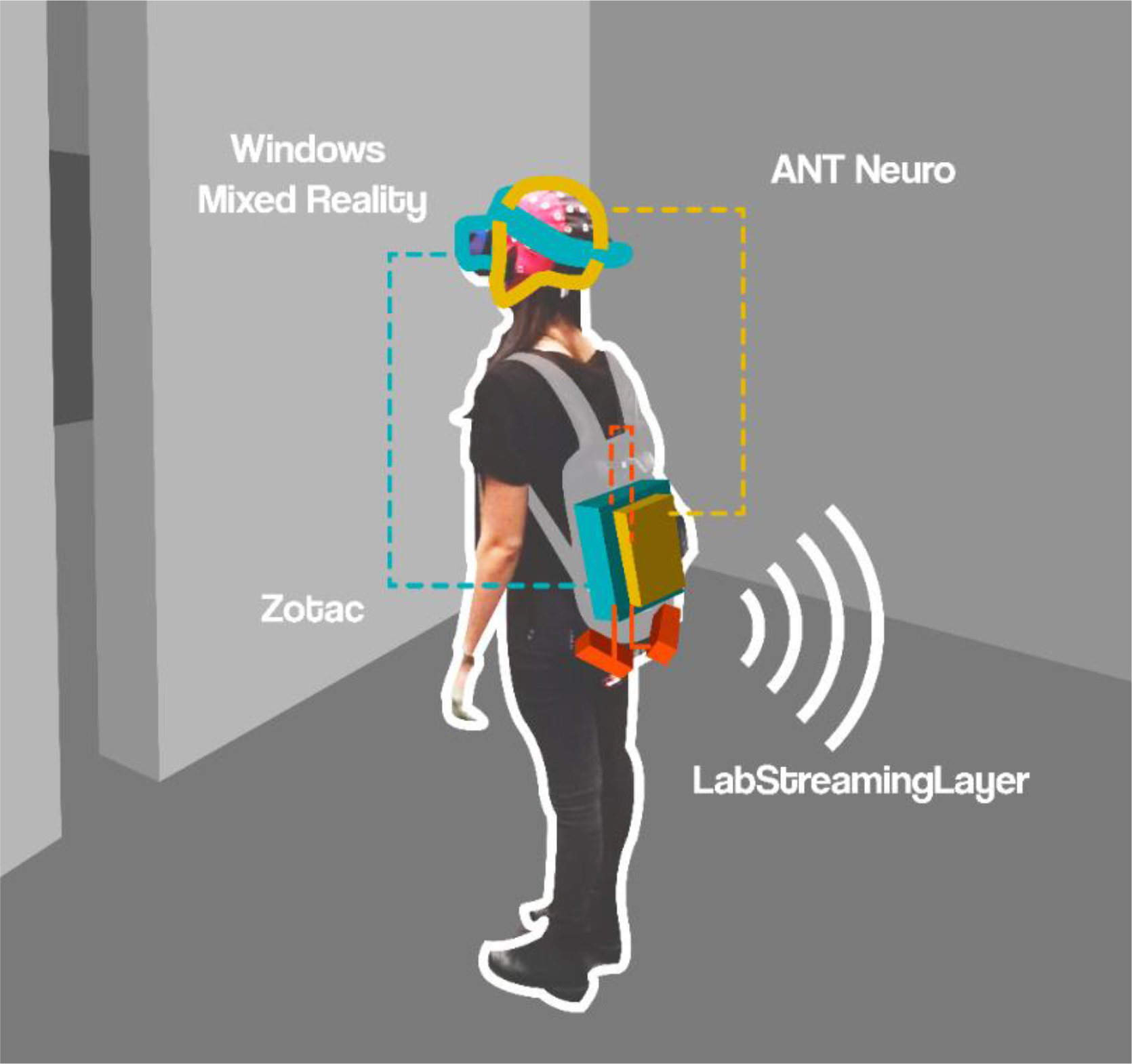
Mobile Brain/Body Imaging setup. The participants wore a backpack, carrying a high-performance gaming computer (Zotac, Cyan color), powered by two batteries (Red color). An EEG amplifier (ANT eegoSports, Yellow color) was attached to the backpack and connected to the computer. The participants wore a VR head mounted display (Windows mixed reality) on top of a 64 channel cap. This setup allowed participants to freely move around while recording data.

EEG data was acquired continuously with a 64 channels EEG system (eegoSports, ANT Neuro, Enschede, Netherlands), sampled with 500 Hz. Impedances were kept below 10 kOhm. The computational delay generated by the interaction of ANT Neuro software, Windows Mixed Reality and Unity was measured to be 20 ms (σ = 4), which was taken into account during the analysis by subtracting the average delay from each event latency. With a jitter of 4 ms, we considered the delay to have little to no impact on the ERPs. Offline analysis were conducted using MATLAB (MathWorks, Natick, MA, USA) and the EEGLAB toolbox (46). The raw data were band-pass filtered between 1 Hz and 100 Hz and down-sampled to 250 Hz. Channels with more than five standard deviations from the joint probability of the recorded electrodes were removed and subsequently interpolated. The datasets were then re-referenced to an average reference and adaptive mixture independent component analysis (AMICA; Palmer et al., 2011) was computed on the remaining rank of the data using one model with online artifact rejection in five iterations. The resultant ICA spheres and weights matrices were transferredtothe raw datasetthat was preprocessed using the identical preprocessing parameters like the ICA dataset, except the filtering, which used a band-pass filter from 0.2 Hz to 40 Hz. Subsequently, independent components (ICs) reflecting eye movements (blinks and horizontal movements) were removed manually based on their topography, their spectrum, and their temporal characteristics.

Epochs were created time-locked to the onset of the room including the closed door (“Lights on”) from −500 ms before to 1500 ms after stimulus onset for *Narrow*, *Mid* and *Wide* door trials. Similarly, another set of epochs were time-locked to the second stimulus *Go/NoGo* from −500 ms before to 1000 ms after onset of the stimulus for *Narrow*, *Mid* and *Wide* door trials. On average, 15% (σ = 10.8) of all epochs were automatically rejected when they deviated more than five standard deviations from the joint probability and distribution of the activity of all recorded electrodes.

The visual-evoked potentials as well as MRCPs were analyzed at central midline electrodes (*Fz*, *FCz*, *Cz*, *Pz*, *POz* and *Oz*) covering all relevant locations including the visual and the motor cortex as reported in previous studies (39, 48). As stimuli were distributed across the complete visual field and participants walked through the virtual spaces, we did not expect any lateralization of ERPs. All channels were analyzed, however only three channels (*FCz*, *Pz* and *Oz*) are reported and discussed in-text according to reported results by Bozzacchi etal. (39). The analysis results of all six channels can be found in the supplementary material. For peak analysis of the P1-N1 complex, the grand average peaks were estimated and individual peaks were defined as the maximum positive and negative peak in the time window surrounding the grand average P1 and N1 peak (+/− 10 ms from peak), respectively. An automatic peak detection algorithm detected the peaks in the averaged epochs for each participant. Multiple peaks were detected and systematically weighed depending on the magnitude, the distance to the grand-average peak latency that was determined by visual inspection of grand average ERP, and the polarity (please see algorithm in the supplementary material). For anterior N1 and posterior P1, by visual inspection of the grand average ERPs, the grand-average latency was estimated to be 140 ms with a search window for individual peaks ranging from 50 - 200 ms. For the anterior P1 and posterior N1 the grand-average peak latency was estimated to 215 ms with a search window for individual peaks ranging from 140 - 290 ms.

Mean peak amplitudes were analyzed using a 3 × 3 repeated measures ANOVA using the door width (*Narrow, Mid, Wide*) and electrode as repeated measures. The results descriptions focus on the visual evoked P1 component at posterior electrodes (*Pz, POz* and *Oz*) and the N1 component at frontal leads (*Fz, FCz* and *Cz*) basedon separate ANOVAs. For the N2 and P2 component at posterior electrodes (*Pz, POz* and *Oz*) and frontal leads (*Fz, FCz* and *Cz*), separate ANOVAs were computed in the time-range of 140 - 290 ms. For the later motor related potentials, an ANOVA was computed for the mean amplitude in the time-range from 600 to 800 ms. The data was analyzed using a 2 × 3 × 6 factorial repeated measures ANOVA with the factors imperative stimulus (*Go* and *NoGo*), door width (*Narrow*, *Mid* and *Wide*), time window (600-700 ms, 700-800 ms) and electrode location (*Fz, FCz, Cz, Pz, POz* and *Oz*). For post-hoc analysis, the data was contrasted using Tukey HSD. In case of violations of the sphericity, corrected p-values are reported. All ANOVA were computed as linear mixed models and all p-values for Tukey HSD contrasts were adjusted using Bonferroni method to account for “within” study design.

## RESULTS

### Subjective and Behavioral results

#### SAM Ratings

A 2 × 3 factorial repeated measures ANOVA with the factors imperative stimulus (*Go* and *NoGo)* and door width (*Narrow, Mid* and *Wide*) for each emotional dimension of the SAM questionnaire revealed differences in the main effect for width: *Arousal* (*F*_*2,4326*_ = *95.12, p* < *0.0001*), *Dominance* (*F*_*2,4326*_ = *46.42, p* < *0.0001*) and *Valence* (*F*_*2,4326*_ = *188.65, p* < *0.0001*). For the imperative stimulus, differences were found for *Arousal* (*F*_*2, 4326*_ = *443.54, p* < *0.0001*), *Dominance* (*F*_*2, 4326*_ = *435.49, p* < *0.0001*), and *Valence* (*F*_*2, 4326*_ = *446.20, p* < *0.0001*). Interaction effects revealed significant difference for all interactions (*all p < 0.0001*). Post-hoc contrasts using Tukey HSD (Figure 3) showed no significant differences for *NoGo* in *Arousal*, however significant differences were identified for *Go* between *Narrow-Mid* (*p < 0.0001*), *Narrow-Wide* (*p < 0.0001*) and *Mid-Wide* (*p < 0.0001*). For *NoGo* in *Dominance* no significant differences were revealed between *Narrow-Mid* (*p = 0.1376*), as opposed to *Narrow-Wide* (*p < 0.0001*) and *Mid-Wide* (*p = 0.0334*), whereas for *Go* no significant differences were found for *Mid-Wide* (*p = 0.2199*), as opposed to *Narrow-Mid* (*p < 0.0001*) and *Narrow-Wide* (*p < 0.0001*). For *Valence*, significant difference were revealed for all contrasts for *Go, Narrow-Mid* (*p < 0.0001*), *Narrow-Wide* (*p < 0.0001*) and *Mid-Wide* (*p < 0.0001*). However, for *NoGo* significant differences were only identified for *Narrow-Mid* (*p < 0.0001*) and *Narrow-Wide* (*p < 0.0001*).

**Fig. 3.**
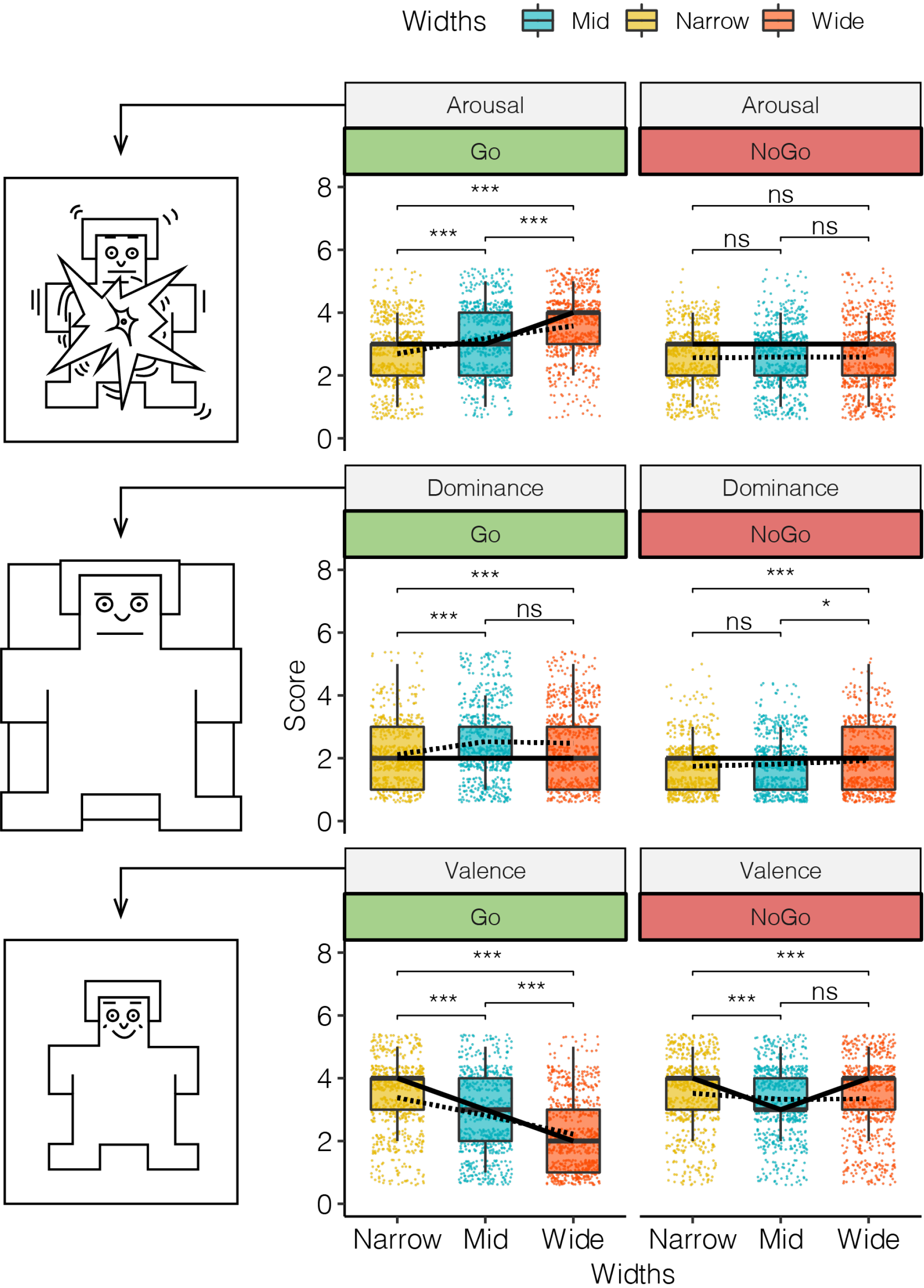
Box plot of the SAM questionnaire results for the three different SAM scales (*arousal*, *dominance*, and *valence*) as a function of the door width (*Narrow*, *Mid*, *Wide*). The left column displays the pictorial representation of the SAM manikin for the highest value of each condition presented. The middle column displays the SAM ratings for the Go condition. The right column displays the SAM ratings for the NoGo-condition. Means are indicated by dashed line, while medians are solid line. Adjusted p values are reported.

##### Performance

To investigate the time it took participants from the *Go*-stimulus to passing the door, a one-way ANOVA with repeated measures for different door widths was computed revealing a significant difference for the factor door widths (*F*_*2,36*_ = *6.404, p* = *0.0042;* Figure 4). Post-hoc comparison (Tukey test) showed no significant differences in behavior when approaching the *Narrow* or *Mid* wide doors (*p > 0.1*), a tendency to be slower when approaching *Mid* as compared to *Wide* doors (*p < 0.1*), and a significant difference between approaching *Narrow* as compared to *Wide* door (*p < 0.001*) with significantly faster approach times for the *Wide* door condition.

**Fig. 4.**
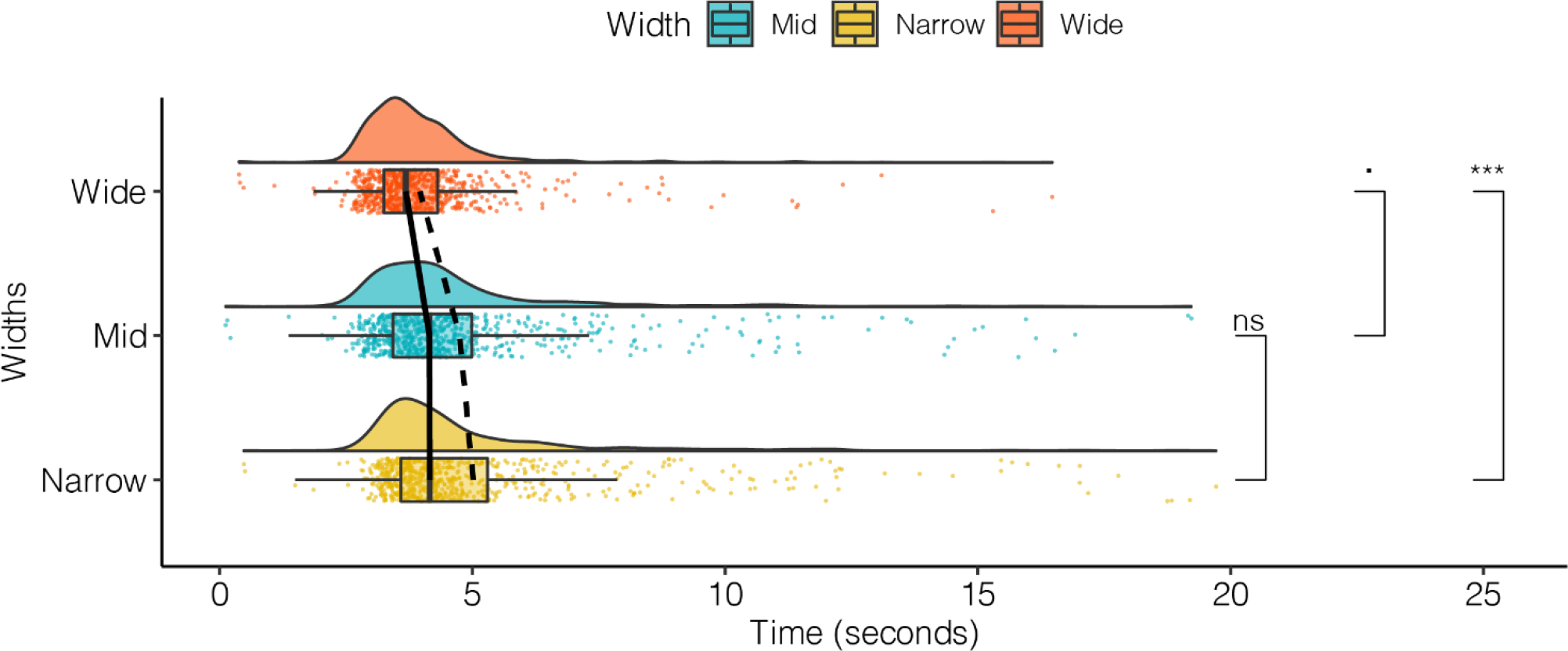
Rain-cloud plot of approach times for each door width condition. Post-hoc comparisons using Tukey test are displayed with a dot < 0.1 and * < 0.05 and *** < 0.001. Means are indicated by dashed line, while medians are displayed as solid lines.

#### EEG - Early event-related potentials

##### Posterior P1

With onset of the lights that allowed participants to see the room including the door (“Lights on”), the ERPs demonstrated a clear P1-N1 complex most pronounced over the occipital midline electrode with a first positive component around 100 ms, followed by a negative peak around 200 ms (Figure 5.1 and see Figure 5.2 in supplementary materials for full six channels). At the frontal midline electrode, this pattern was inversed and a negative component around 100 ms was followed by a positive peak observed around 200 ms. The 3 × 3 repeated measures ANOVA on P1 amplitudes for posterior electrodes revealed significant main effects for both the factors widths (*F*_*2,108*_ = *8.163, p* = *0.005*) and channel (*F*_*2,36*_ = *15.868, p* < *0.0001*). The interaction effect was not significant (*F*_*4,108*_ = *1.669, p* = *0.1624*). Post-hoc comparisons using Tukey HSD test revealed significant differences in peak amplitudes at channel *Oz* between *Narrow* and *Mid* wide transitions (*p = 0.0021*) and between *Narrow* and *Wide* transitions (*p = 0.0065*) but no differences between *Mid* and *Wide* transitions (*p = 1*). Tukey contrasts yielded no significant differences between electrodes, with differences in P1 amplitudes at *POz* comparing *Narrow* and *Wide* transitions (*p = 0.028*).

**Fig. 5.1.**
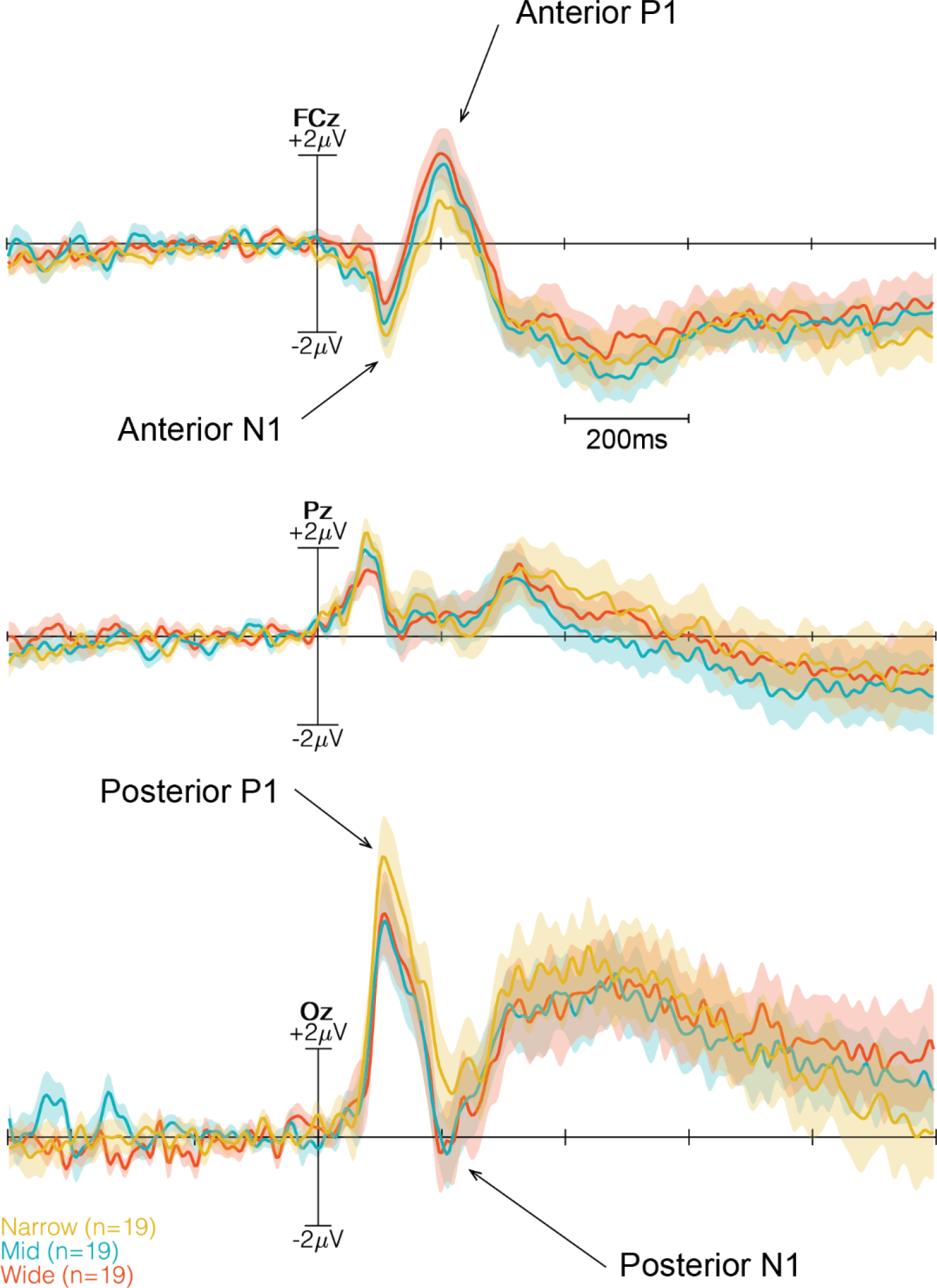
Three time-locked ERPs (*FCz, Pz* and *Oz*) at the onset of “Lights On” event. *Narrow* condition in yellow, *Mid* condition in blue and *Wide* condition in red. Two time windows are indicated with dashed-lines and grey transparent box. The first time window (50 - 200 ms) mark the anterior N1 and posterior P1, while the second window (140 - 290 ms) mark the anterior P1 and posterior N1. The components are marked with arrows.

##### Posterior N1

The 3 × 3 repeated measure ANOVA on N1 amplitudes for posterior electrodes revealed a significant main effect for the factor door widths (*F*_*2,108*_ = *4.348, p* = *0.0153*) and no significant impact for the factor channels (*F*_*2,36*_ = *0.0893, p* = *0.9147*), nor the interaction (*F*_*4,108*_ = *1.304, p* = *0.2731*). Post-hoc Tukey HSD contrasts revealed no significant differences for *Pz* and *POz*. However,similar to posterior P1, significant differences at *Oz* for the comparison of Narrow and Mid wide transitions (*p = 0.0113*) and for the comparison of *Narrow* and *Wide* transitions (*p = 0.0372*) were found (Figure 6).

**Fig. 6.**
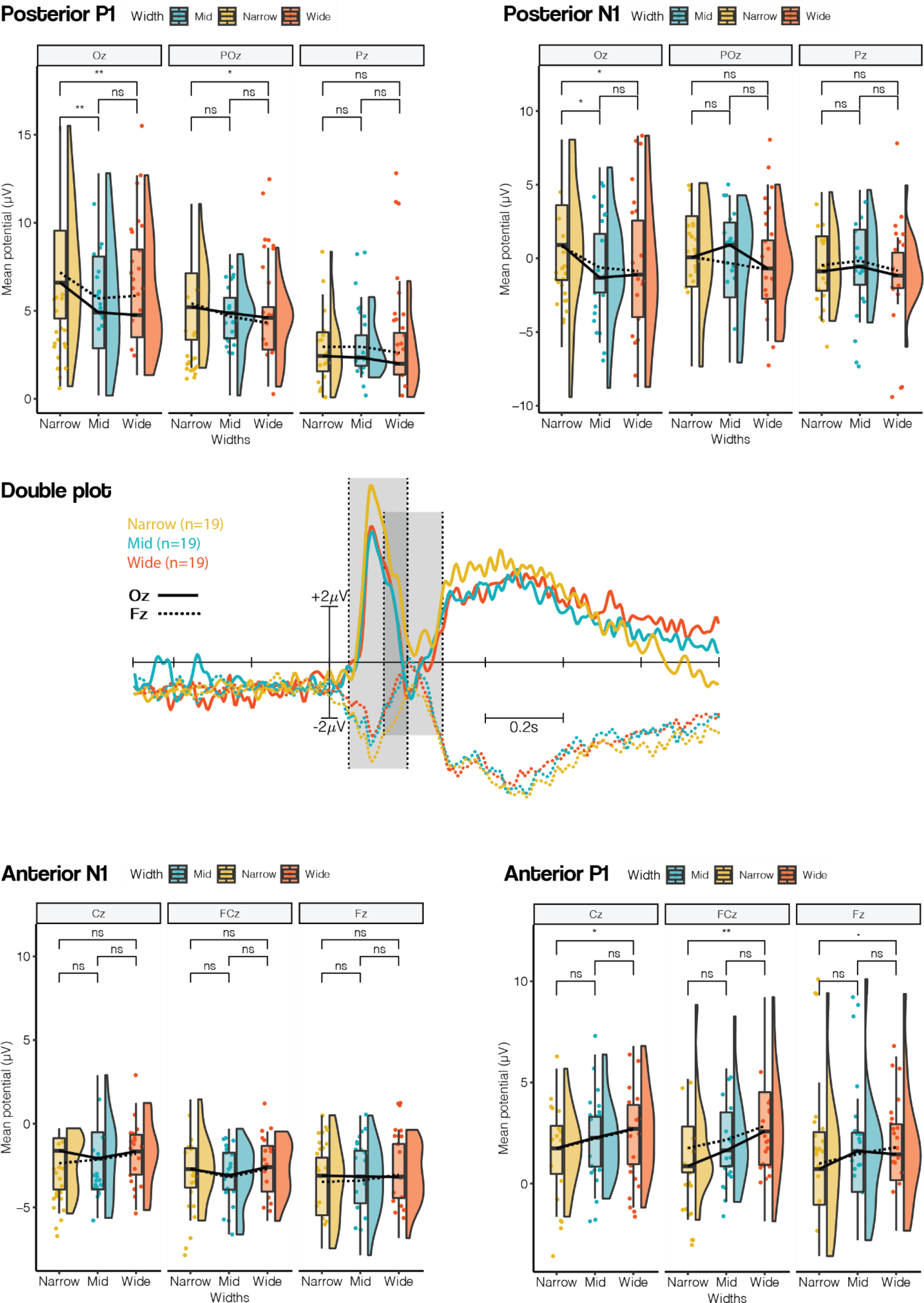
Posterior P1. Rain-cloud plot of detected mean amplitude of positive peak in time-locked event “Lights on” in the time range of 50 to 200 ms for *Pz, POz* and *Oz*. Means are indicated by dashed line, while medians are solid line. Significance is calculated using Tukey HSD. We observed significant differences for *Oz* between *Narrow-Mid* (*p = 0.0021*) *and Narrow-Wide* (*p = 0.0065*), while for *POz* in *Narrow-Wide* revealed significant difference *(p = 0.028)*, however no significant differences were observed in other electrodes and other contrasts. **Posterior N1.** Rain-cloud plot of detected mean amplitude of negative peak in time-locked event “Lights on” in the time range of 140 to 290 ms for *Pz, POz* and *Oz*. We observed significant differences only for *Oz* in *Narrow-Mid* (*p = 0.0113*) and *Narrow-Wide* (*p = 0.0372*). **Anterior N1.** Rain-cloud plot of detected mean amplitude of negative peak in time-locked event “Lights on” in the time range of 50 to 200 ms for *Fz, FCz* and *Cz* We observed no significant differences for any electrode. **Anterior P1.** Rain-cloud plot of detected mean amplitude of negative peak in time-locked event “Lights on” in the time range of 140 to 290 ms for *Fz, FCz* and *Cz* We observed significant differences in all electrodes in *Narrow-Wide*, with the exception of only a tendency in *Fz* (*p = 0.0717*), *FCz* (*p = 0.0071*) and *Cz* (*p = 0.0214*). **Double plot.** Frontal (dashed-line) and posterior (solid-line) time-locked ERPs (*Fz* and *Oz*) at the onset of “Lights On” event. *Narrow* condition in yellow, *Mid* condition in blue and *Wide* condition in red. Two time windows are indicated with dashed-lines and grey transparent box. The first time window (50 - 200 ms) mark the anterior N1 and posterior P1, while the second window (140 - 290 ms) mark the anterior P1 and posterior N1.

##### Anterior P1

An inverse pattern was observed for amplitudes over anterior leads with a main effect of door widths that differed depending on the affordances (*F*_*2,108*_ = *11.071, p* < *0.0001*). The main effect of channels also reached significance (*F*_*2,36*_ = *5.3627, p* = *0.0092*). Tukey HSD contrasts revealed significant differences only between *Narrow* and *Wide* transitions for *FCz* (*p = 0.0071*) and *Cz* (*p = 0.0214*), and a tendency at *Fz* (*p = 0.0717*). The interaction was not significant.

##### Anterior N1

The 3 × 3 repeated measures ANOVA on N1 amplitudes for anterior electrodes revealed no significant main effect for the factor door widths (*F*_*2,108*_ = *1.823, p* = *0.1663*). In contrast, the main effect of channels reached significance (*F*_*2,108*_ = *8.109, p* = *0.0012*). The interaction did not reach significance.

##### EEG - Motor-related processes

After onset of the imperative stimulus a positive peak at anterior leads and a negative peak at posterior leads were observed. For sake of brevity, this potential complex is referred to as early post imperative complex (EPIC). Reflecting similar cortical polarity as the P1-N1 complex, the EPIC was analyzed in a similar way, separating anterior leads (*Fz, FCz* and *Cz*) from posterior leads (*Pz, POz* and *Oz*), and detecting single peaks in individual averages.

##### Anterior EPIC

A 2 × 3 × 3 repeated measures ANOVA revealed significant difference in the main effect for widths (*F*_*2,270*_ = *4.21, p* = *0.0157*), imperative stimulus (*F*_*1,270*_ = *23.66, p* < *0.0001*), and for channel (*F*_*2,36*_ = *6.70, p* = *0.0033*). Nointeraction effectwas observed. The Bonferroni-corrected post-hoc Tukey HSD revealed no significant differences between the transition widths for different channels or imperative stimuli.

##### Posterior EPIC

The identical ANOVA for the posterior potentials of the EPIC revealed no significant impact of transition widths (*F*_*2,270*_ = *2.001, p* = *0.1371*) nor imperative stimulus (*F*_*1,270*_ = *2.30, p* = *0.1298*). Significant differences in EPIC amplitude were observed for the factor channel (*F*_*2,36*_ = *5.45, p* = *0.0085*). Since topographical differences were not in the focus of this study, no further post-hoc contrasts were computed. No interaction was significant.

##### PINV

In the preparation time prior to the onset of the door color change, indicating either to walk through the door or to remain in the same room, we observed no systematic negative going waveform as reported in previous studies (37, 49). However, after onset of the color change, a pronounced positivity, followed by a long-lasting negative waveform over fronto-central locations was observed in the ERP (Figure 7.1 and see Figure 7.2 in supplementary material for full six channels). This negative waveform resembled a post-imperative negative variation (PINV) as described in previous studies (40, 42, 50). The PINV component was observed 600-800 ms post imperative stimulus (color change of the door) and varied as a function of the affordance of the environment (door width). A global 2 × 3 × 6 factorial repeated measures ANOVA was computed to analyze the MRCPs using *Go/NoGo*, *Width* and *Electrode* as repeated measures. The ANOVA revealed significant differences in the main effect for *Go/NoGo* (*F*_*1,540*_ = *19.54, p* < *0.0001*) and for *Electrode* (*F*_*5,90*_ = *16.69, p* < *0.0001*). Significant differences were reported for the interaction effect of *Go/NoGo:Channel* (*F*_*5,540*_ = *5.25, p* = *0.0001*) and for *Width:Channel* (*F*_*10,540*_ = *2.61, p* = *0.0042*).

**Fig. 7.1.**
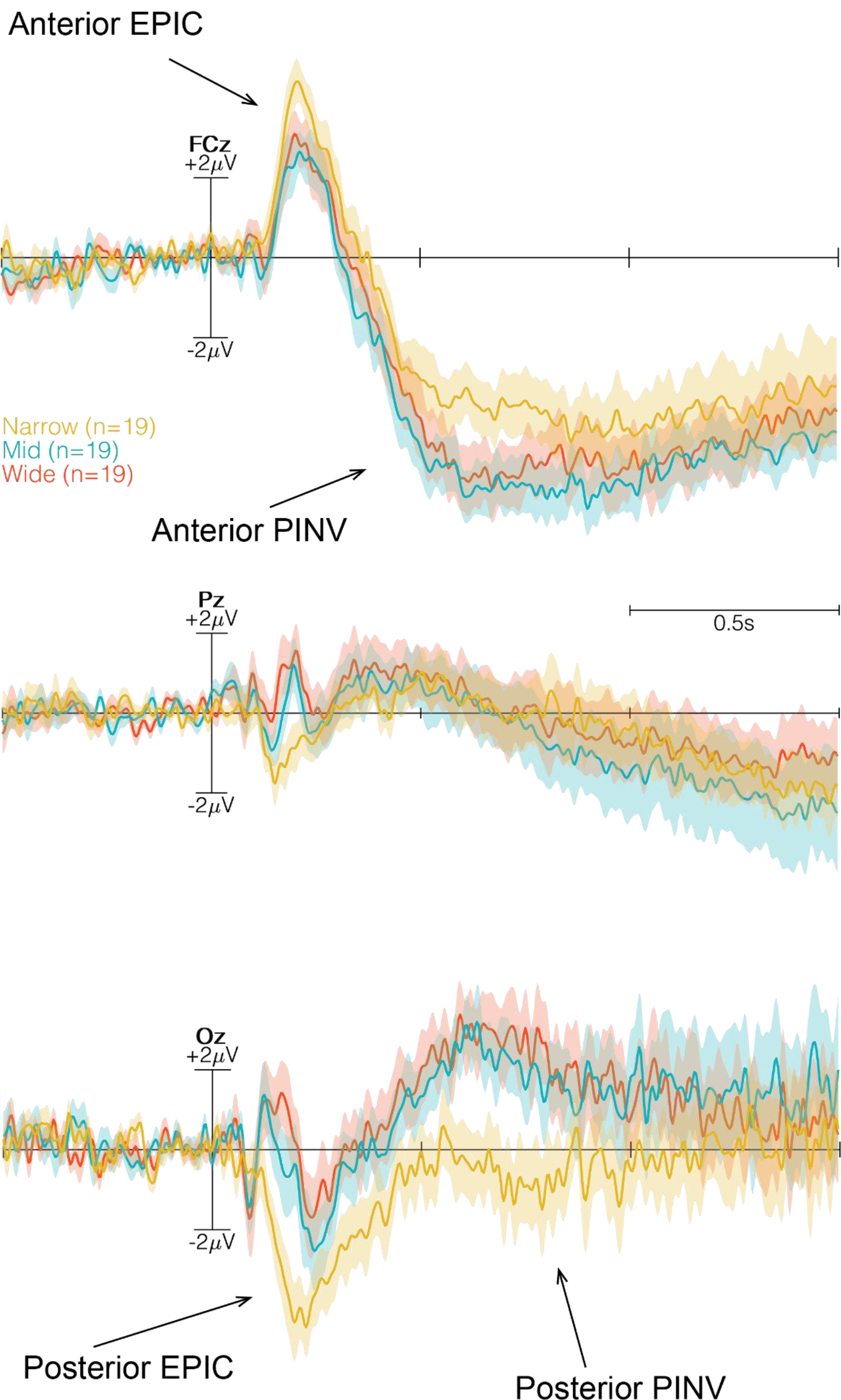
Three time-locked ERPs (*FCz, Pz* and *Oz*) at the onset of Go/NoGo. *Narrow* condition in yellow, *Mid* condition in blue and *Wide* condition in red. The time window, indicated with dashed-lines and grey transparent box, illustrates the selected time window to analyze the MRCP by a global 2 × 3 × 6 factorial repeated measures ANOVA. Anterior and posterior PINV are marked with arrows.

Post-hoc contrasts, using Tukey HSD, revealed significant differences only for the *Go* condition, as opposed to the *NoGo* condition (Figure 8.1). Similar to the early evoked potentials, differences were only observed in frontal and occipital sites and between *Narrow* and *Mid* door widths over *FCz* (*p = 0.0059*) and *Oz* (*p < 0.0001*), as well as between *Narrow* and *Wide* doors at *FCz* (*p = 0.0323*) and *Oz* (*p < 0.0001*). No differences were observed betweenthe *Mid* and *Wide* doors (Figure 8.2 in the supplementary material for all six channels).

**Fig. 8.1.**
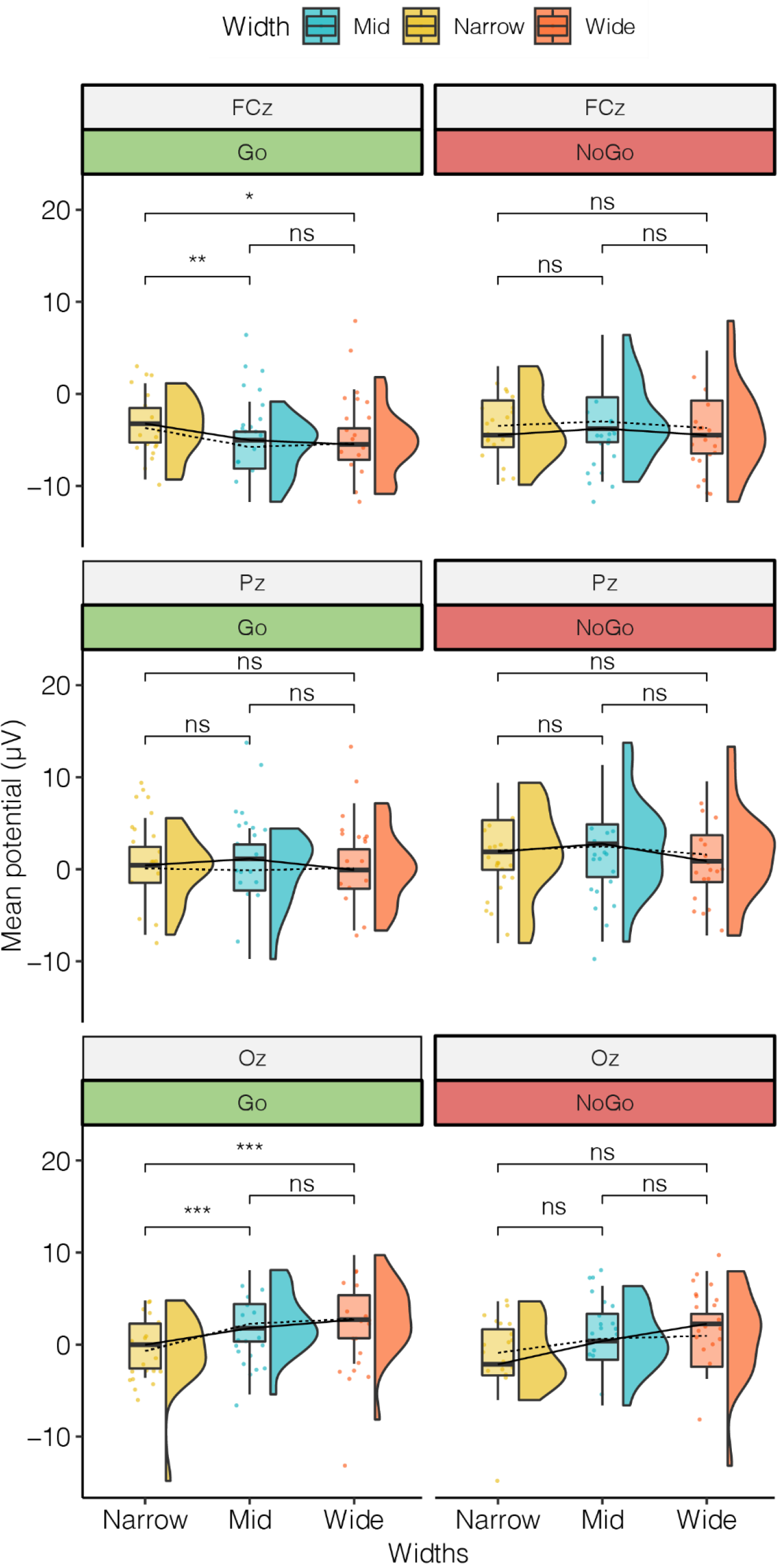
Rain-cloud plots of mean amplitude of negative development in time-locked event of Go/NoGo in the time range of 600 to 800 ms for *FCz, Pz* and *Oz*. Means are indicated by dashed line, while medians are solid line. The Tukey HSD contrast revealed differences only in *FCz* and *Oz*, and between *Narrow-Mid* for *FCz* (*p = 0.0059*) and for *Oz* (*p < 0.0001*), and between *Narrow-Wide* for *FCz* (*p = 0.0323*) and for *Oz* (*p < 0.0001*). No differences were observed for NoGo.

### DISCUSSION

The main goal of this study was to assess whether brain activity is altered depending on the affordances offered by the environment. If such an account holds true, affordances should systematically modulate behavior and brain activity. Specifically, we hypothesized that perceptual processes co-vary with the environmental affordances leading to behavioral changes and that motor-related cortical potentials would vary as a function of affordances.

#### SAM and approach time

The results of the questionnaire should be interpreted with caution due to the amount of trials per participant, the varying sensitivity to VR and the different skills of subjective emotional evaluation. The analysis of subjective ratings revealed significant differences between different *Go* trials, but no differences for *NoGo* trials regarding *Arousal* ratings. When given a *NoGo*, participants responded perhaps arbitrarily, feeling unburdened, causing no significant difference among the three door widths. Notably, in cases of *NoGo*, all participants perceived a similar scene standing in front of a red (*NoGo*) door, turning around and answering the virtual SAM. The only variable in this sense was the door width, while the only difference from *NoGo* to *Go*, was the action itself. The subjective ratings highlight the influence of action on evaluating the environment. If space was to be investigated statically (comparable to the case of *NoGo*), we would not have been able to detect any differences for *Arousal* for varying door sizes, potentially due to the absence of action. Varying door sizes for *Go* trials yielded differences between passable and impassable conditions for *Dominance*, reporting that *Narrow* door was more dominating than *Mid* and *Wide*. However, for *Valence* we observed an increasing score the narrower the door, which is the opposite behavior observed for *Arousal*. These results indicate that being able to pass easily is more exciting, less pleasant and less dominating. This effect is perhaps grounded in the monetary reward participants could receive only when successfully passing through to the next room. Most importantly, however, the findings indicate that subjective reports differ significantly dependent on whether participants actively moved through the rooms or not implying an impact of action affective ratings of an environment. We speculate whether the omnipresent significant differences may be rooted in uniqueness of emotional states that varies from participant to participant. Such an account of emotional ratings is currently gaining credibility (51, 52).

The time it took participants to reach the door after onset of the imperative color change varied according to the environmental affordance. Participants approached the *Wide* door significantly faster than *Mid* and *Narrow* doors, while there was no significant difference for *Mid* and *Narrow* transitions. While the *Wide* door clearly offered a passage without greater computational demands_regarding the motor plan and execution, the *Mid* door width, being ambiguously wide/narrow, might have triggered motor processes simulating a transition to estimate whether the door was passable or not. In this sense, the *Mid* and *Narrow* doors, causing uncertainty, might have delayed approach times due to increasing processing demands. Admittedly, results derived from the approach time are limited, partly due to the caused fatigue of operating a physically demanding task for a relatively long time period, and partly due to the subjective manner and interpretation of passing a door that is seemingly impossible to pass. This caused participants to develop different approach strategies which caused different delays. However, the fact that participants, in general, spent significantly more time approaching the *Narrow* doors compared to *Wide* doors provides sufficient guidance for the analyses of cortical measures associated with these differences.

#### Cortical measures

##### Early evoked potentials

As an initial insight into the association of affordances and cortical potentials, we analyzed the early visual-evoked potentials. We expected to find differences in the stimulus-locked ERP at occipital channels reflecting differences in sensory processing of affordance-related aspects of the transition. Importantly, based on the assumption of fast sensorimotor active inferences that should be reflected in action-directed stimulus processing influencing not only sensory but also motor-related activity, we hypothesized to also find differences in the ERP over motor areas in the same time window as sensory potentials (i.e., between 50 and 200 ms). As illustrated in the analysis, we found significant differences in amplitudes of the visually evoked P1 component over the central occipital electrode dependent on the affordance of the transition. In addition, in line with our hypothesis, we also found a difference over fronto-central leads starting around 50 ms and lasting until 200 ms after onset of the doors display. Taken together, no significant differences in peak amplitudes were found when comparing the passable *Mid* and *Wide* doors while peak amplitude associated with both door widths significantly differed from impassable *Narrow* doors. Note that the visual scene of the three doors are comparable as they contained same physical contrasts, and that participants at this point did not know whether to go or not as they were merely introduced to the setting they might have to pass in a couple of seconds. As no significant differences were found for *NoGo*, it functions as a matching control, and thus we can interpret the differences in *Go* as affordance manipulation. These results indicate that impassable doors with poor affordances produce significantly different early evoked potentials compared to passable doors particularly at fronto-central and occipital sites. Thus, environmental affordances, in terms of being able to program bodily trajectory to transit spaces, yield a significant measurable effect on early cortical potentials best pronounced over frontal and occipital sites at approximately 200ms after first view of the environment.

Considering the affordance-specific pattern observed for the early P1-N1-complex, prior studies have shown this visual evoked potential complex to reflect attentional processes associated with spatial or feature-based aspects of stimuli (53–57). Attended stimuli elicit larger P1-N1 amplitudes than unattended ones. Based on these findings, the results suggest that passable transitions were associated with increased attentional processing. Approaching the affordance-specific pattern of P1-N1-complex using active inferences (58), the difference confirms the assumption that perceptual processes co-vary with environmental affordances. In this sense, the amplitude difference might be credited to the process of active inference of whether the body can actively move and transit at all. This implies that visual attention is also guided by action-related properties of the environment and support the concept of fast, lower sensorimotor active inferences, explained as hierarchical and dynamic model of the world. Similar to HAC (31) and active inference (30, 59), these findings are in line with parallel cortical processes integrating sensory information to specify currently available affordances. Similarly, this means that, how one might act upon the environment is an ongoing process of affordances, taking place as early as perceptual processes, and which situates actions in an intimate position with perception. Such early processes are deeply involved in the impression of the environment for an agent pointing towards the importance of movement in cognition, and of how an agent enacts the world. Given affordances are processed at such an early stage, we speculate whether the impression of an environment compose the immediate experience of the environment in a particular setting. Such an immediate experience fits with the term *atmospheres* as defined by Zumthor (60) “I enter a building, see a room, and - in a fraction of a second - have this feeling about it”, and thus relating the instantaneous emerging experience of space to affordances and action in general.

##### Motor-related potentials

Although the ERP plots indicate an affordance-trend of the EPIC, statistical tests revealed no significant differences. However, *Narrow* door width elicited the greatest amplitude, both in case of anterior positivity and posterior negativity. In line with prediction errors and affordances, the increased amplitude associated with *Narrow* transitions can be interpret as a reflection of the body simply not fitting, and yet forced to interact with the transition. Recall that prior to the imperative stimulus, participants have been standing for 6 s (σ = 1 s). The EPIC may have an influence on the PINV. The nature of the PINV component is not as well investigated as other ERP components, limiting the reliability of an interpretation based on only a few studies that treat the component as modality-unspecific, and rather “*consider the PINV as an electrocortical correlate of a cognitive state”* (61). Since the study by Gauthier and Gottesmann (62) the PINV, similar to affordances, has been hypothesized to act as a marker of change in psychophysiological state. Ever since, the PINV has been used to investigate depression, schizophrenia, learned helplessness and loss of control (40–42, 63, 64). Results show depressive and schizophrenic participants to exhibit an increased PINV that is explained as increased vulnerability for loss of control, as well as increased anticipation for future affective events (40, 42, 50). If an increased PINV reflects increased vulnerability for future events, as we observed for impassable doors, then the component, constituted by continuous motor potential activity, sheds new light on affordances as an intrinsic affective property of action itself. Casement and colleagues (42) even suggested the PINV to depend on lack of control as the state of having no influence; depriving the potential to act. This could explain the difference in the *Narrow* condition, as participants were instructed to attempt to pass at all times until failure leading to a sense of loss of control. Only in cases of *Go* did we observe a difference in the PINV component, which varied similar to the P1-N1-complex. Amplitudes of the component for *Narrow* doors were significantly different from *Mid* and *Wide* doors, while the passable conditions did not differ from one another. Further, there were no significant differences in the PINV component in cases of *NoGo*, emphasizing the importance of the motor execution itself to evoke the PINV component. These results point towards the PINV component as an expression of willingness to execute an act restricted beyond ones’ own control, i.e. a designed environment. Thus, the PINV might serve as an excellent marker for affordances.

The presented results of the PINV are consistent with the observed increase in activity over fronto-central sites by Bozzacchi et al. (39). Bozzacchi and colleagues concluded that the meaning of the action and awareness of being able to act - affordances - affect action preparation, which is here understood as the motor-related potential prior to movement onset. We argue that the PINV component might reflect a willingness, or even intentional, aspect of affordances. This would mean that the PINV is not modulated by the perception (that the door is a different visual information), but reveals something about the intention of movement - which we translate to affordances. For this reason, we find significant differences in cases of *Go*, but not in *NoGo*, and further for passable compared to impassable. In light of HAC (31), a potential explanation for the absence of differences in the *NoGo* trials, is related to the immediate action selection, which in all cases (*Narrow, Mid* and *Wide*) is a simple turn to answer the questionnaire, and thus present the participant with identical affordances. When instead given a *Go*, cortical processes require an action selection related to the anticipated motor trajectory, which differs according to the affordances of the door width. Regarding the temporal aspect of transitioning to the next room, HAC suggests the higher levels bias the lower level competitions, which operate at the level of action itself, through a cascade of expected next affordances. The lower levels have a continuous competition of how to satisfy the higher expectations. Action selection, executed while unfolding the planned movements in a continuous manner, depend on the expectation of next affordances. Taken together, the post-hoc analyses revealed differences grouped for passable as compared to impassable doors throughout all channels, except for *Pz*. We do not observe any differences between *Mid*-*Wide*, but find significant differences between *Narrow-Mid* and *Narrow*-*Wide*. The greatest differences were found over fronto-central and occipital sites. Similar to the early evoked potentials, these results indicate that environmental affordances impact neural activity prior to action depending on whether one has to act or not.

Notably, regarding architectural experience, since the PINV component was only expressed in the *Go* condition (forced interaction with the environment), these findings support the importance of movement for architectural experience, in a sense that action or even only the perception of action possibilities alters brain activity. Visually guiding and propelling the body in space greatly influences the continuous emerging of affordances, which in turn affect the human experience. We found differences in fronto-central and occipital areas, prior to movement through space with the post-imperative negative going waveform most pronounced over *FCz* indicated an involvement of the supplementary motor area (SMA) as reported by Bozzacchi et al. (39). Interestingly, earlier studies showed involvement of SMA in visually guided actions (65), which is the essence of active inferences. The PINV can be generated independently from the re-afferent signal, which is, in terms of active inference, understood as ascending (bottom-up) proprioceptive prediction-errors (66). This suggests the PINV component might reflect descending (top-down) predictions, rendering SMA as an essential area of action-perception loop, and thus crucial for processing continuous affordances. This account might resolve the finding of fronto-central differences in *Go* trials only. The SMA is anatomically bridging the frontal cortex with motor cortex - perhaps also functionally as argued by Adams et al. (66), as this anatomical nature fits with the proposed hierarchical characteristics of forward and backward projections in active inferences.

### CONCLUSION

The present study provides strong evidence for affordances to be processed as early as perceptual processes, linking action and perception in a similar manner to active inference. The results points towards a conception of the brain that seems to deal with “how can I act” while in parallel processes referringto “what do I perceive” take place. The results thus support the assumption that perception of the environment is influenced by affordances and action itself - hence, affordances and action can influence experience of an environment. Due to the importance of affordances and action for brain dynamics, this further emphasizes and qualifies the general idea of enactivism as a holistic approach to investigate cognition. We do not claim that architectural affordances are directly represented as a specific event-related potential component; however, we provide evidence for an action-perception account of cognition, which systematically differentiates according to the definition of affordances.

The nature of the analyzed brain activity emphasizes the importance of the intentional movement. Our results are consistent with the concept of continuous affordances as explained by active inferences. In terms of architecture, the results shed light on why transitions have been a constant throughout the history of architecture, perhaps especially in religious and other buildings that actively aimed at producing a certain experience of presence. Thus, the fact that we are predictive beings, in terms of architecture, means we should take into consideration how bodily movement alters perception. By altering perception, this would ultimately lead spaces to have a potentially physiological impact on users. Much remains to be uncovered in architectural cognition. Moving and transitioning in space, is continuously constructing a prediction of a world, a world that we perceive dependent on our action potentials, which informs brain, body and mind. Transitions in architecture form a holistic entity of architectural experience expressed as the unfolding of motor planning, spatial sequences and predictive mechanisms. Similar to Zeki (67), we speculate whether the ancient interest in tailoring transitions and sequences may have developed as a trial-and-error of active-narration, perhaps rooted in ancient knowledge of the predictive mind and action-perception parallel processing nature of the human being.

## ACKNOWLEDGEMENTS

We thank Lukas Gehrke, Marius Klug and Federica Nenna for their support with the technical setup. We thank in particular Federica Nenna for her additional support in conducting the experiment. In addition, we would like to thank Andrea Jelic for important discussions and bringing essential insights of enactivism to our knowledge.

## COMPETING INTERESTS

The authors declare no competing interests.

## Supplementary figure legends

**Figure 5.2 - supplementary.**
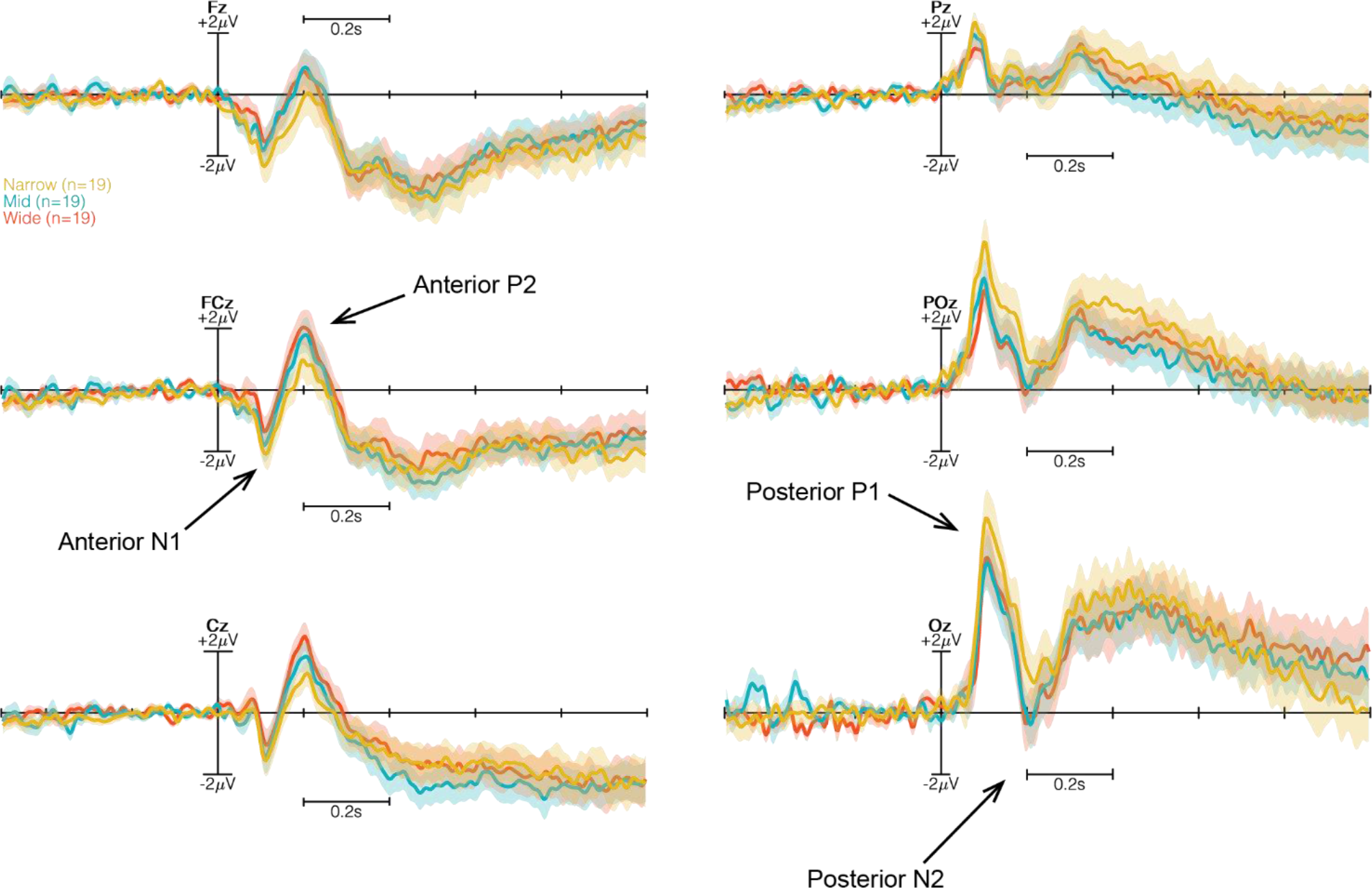
ERP plots of “Lights On” stimulus for all six channels (*Fz, FCz, Cz, Pz, POz* and *Oz*). *Narrow* condition in yellow, *Mid* condition in blue and *Wide* condition in red. N1-P1-complex are marked with arrows.

**Figure 7.2 - supplementary.**
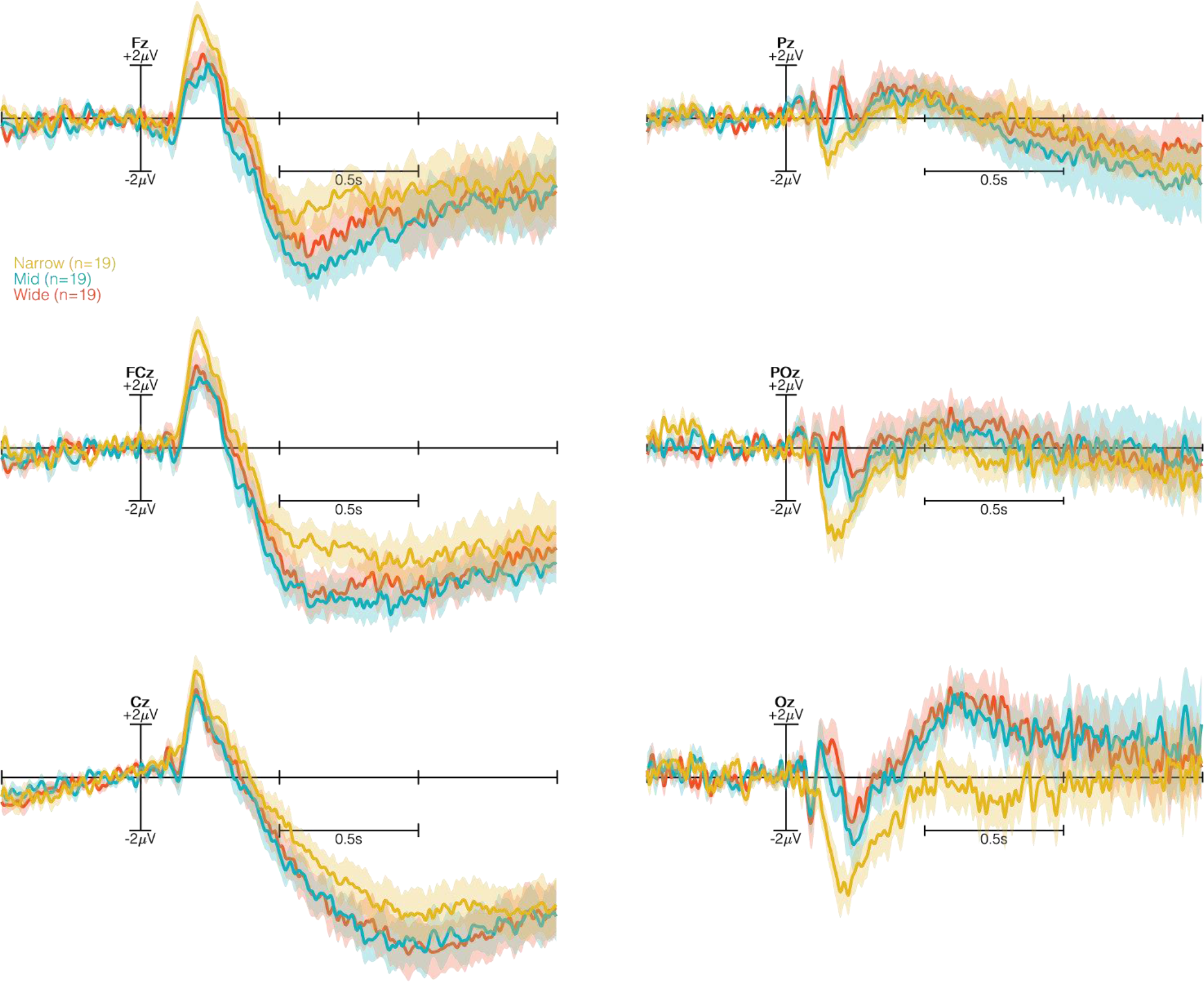
ERP plots of the total six channels only for Go trials. ANOVA with repeated measures of time-locked ERP, where the increasing darkness behind the plots indicates the increasing level of significance. The repeated measures ANOVA revealed Fz (*F*_*2,36*_ = *4.546, p* = *0.0174*), FCz (*F*_*2,36*_ = *7.116, p* = *0.0025*), Cz (*F*_*2,36*_ = *4.116, p* = *0.0236*), Pz (*F*_*2,36*_ = *0.089, p* = *0.915*), POz (*F*_*2,36*_ = *1.708, p* = *0.196*) and Oz (*F*_*2,36*_ = *14.39, p* < *0.0001*). We observed no difference for NoGo - however, we observed a difference within frontocentral and occipital sites for Go trials.

**Figure 8.2 - supplementary.**
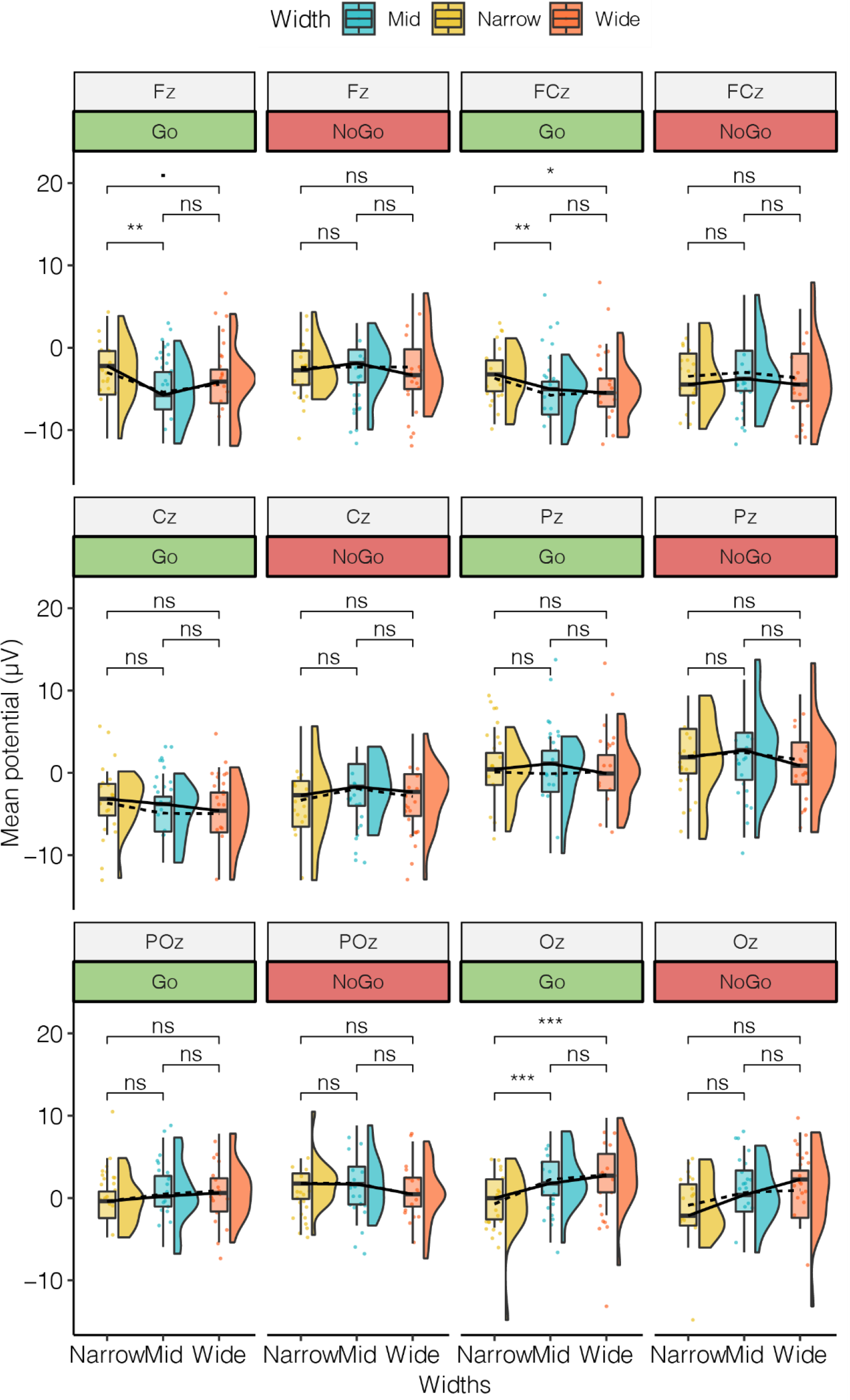
Rain-cloud plot of the mean amplitude of selected six channels between 600 - 800 ms post imperative stimulus - PINV component. Means are indicated by dashed line, while medians are solid line. We compared (Tukey test) the *Width* within Go and NoGo conditions, and observed only significant differences for Go condition. We observed differences within frontocentral and occipital sites.

## REFERENCES

1. Gibson J The Ecological Approach to Visual Perception. Hought Mifflin-Bost, (1979).

2. Clark A An embodied cognitive science? Trends Cogn Sci 3(9):345–351, (1999).

3. Clark A Surfing Uncertainty: Prediction, Action and the embodied mind (Oxford University Press, New York) (2015).

4. Friston KJ, Kilner J Harrison L A free energy principle for the brain. J Physiol 100(1-3):70–87, (2006).

5. Hohwy J The Predictive Mind (Oxford University Press) doi:10.1093/acprof:oso/9780199682737.001.0001 (2013).

6. Friston K The free-energy principle: a unified brain theory? Nat Rev Neurosci 11(2):127–138, (2010).

7. Friston K The free-energy principle: a rough guide to the brain? Trends Cogn Sci 13(7):293–301, (2009).

8. Friston K, Mattout J Kilner J Action understanding and active inference. Biol Cybern 104(1-2):137–160, (2011).

9. Friston K Learning and inference in the brain. Neural Networks 16(9):1325–1352, (2003).

10. Clark A Whatever next? Predictive brains, situated agents, and the future of cognitive science. Behav Brain Sci 36(3):181–204, (2013).

11. OED Oxford English Dictionary (Oxford University Press, Oxford) (2018).

12. Fazio MW, Moffett M Wodehouse L A world history of architecture (Laurence King). 2nd Ed, (2008).

13. Norberg-Schulz C Intentions in architecture. (MIT Pr) (1965).

14. Pallasmaa J The Embodied Image: Imagination and Imagery in Architecture (Wiley) (2011)‥

15. Palladio A The four books on architecture (MIT Press) Available at: https://www.saxo.com/dk/the-four-books-on-architecture_andrea-palladio_paperback_9780262661331 [Accessed October 20, 2018] (1997).

16. Vitruvius, Morgan MH (Morris H Vitruvius : the ten books on architecture (Dover Publications) Available at: https://www.saxo.com/dk/on-architecture-bks-i-x_vitruvius_paperback_9780486206455 [Accessed October 20, 2018] (1960).

17. Rasmussen SE Experiencing architecture (Chapman & Hall, London) (1959).

18. Moretti L, Bucci F Mulazzani M DeConciliis M Luigi Moretti: Works and writings (Princeton Architectural Press) (2002).

19. Arnheim R The Dynamics of Architectural Form (University of California Press) (1977).

20. Boettger T Threshold Spaces : transitions in architecture analysis and design tools (Birkhäuser, Weimar) (2014).

21. Jelić A, Tieri G De Matteis F Babiloni F Vecchiato G The Enactive Approach to Architectural Experience: A Neurophysiological Perspective on Embodiment, Motivation, and Affordances. Front Psychol 7:481, (2016).

22. Holl S, Pallasmaa J Pérez-Gómez A Questions of perception : phenomenology of architecture (William Stout Publishers, San Fransisco) Available at: http://catalog.hathitrust.org/Record/008231335 (2006).

23. Bachelard G The poetics of space (Beacon Press) (1969).

24. Norberg-Schulz C Nightlands (MIT Press, Cambridge, Mass. ; London) (1997).

25. Coburn A, Vartanian O Chatterjee A Buildings, Beauty, and the Brain: A Neuroscience of Architectural Experience. J Cogn Neurosci 29(9):1521–1531, (2017).

26. Varela FJ, Thompson E Rosch E The embodied mind: cognitive science and human experience Available at: https://mitpress.mit.edu/books/embodied-mind-0 [Accessed December 6, 2017] (1991).

27. Thompson E Mind in life: biology, phenomenology, and the sciences of mind (Belknap Press of Harvard University Press) Available at: http://www.hup.harvard.edu/catalog.php?isbn=9780674057517 [Accessed December 6, 2017] (2007).

28. Maturana HR, Varela FJ The tree of knowledge : the biological roots of human understanding (Shambhala) (1992).

29. Gallagher S Enactivist interventions: rethinking the mind (Oxford University Press). 1st Ed, (2017).

30. Kiebel SJ, Daunizeau J Friston KJ A Hierarchy of Time-Scales and the Brain. PLoS Comput Biol 4(11):e1000209, (2008).

31. Pezzulo G, Cisek P Navigating the Affordance Landscape: Feedback Control as a Process Model of Behavior and Cognition. Trends Cogn Sci 20(6):414–424, (2016).

32. Makeig S, Gramann K Jung T-P, Sejnowski TJ Poizner H Linking brain, mind and behavior. Int J Psychophysiol 73(2):95–100, (2009).

33. Gramann K, et al. Cognition in action: imaging brain/body dynamics in mobile humans. Rev Neurosci 22(6):593–608, (2011).

34. Gramann K, Jung T-P, Ferris DP Lin C-T, Makeig S Toward a new cognitive neuroscience: modeling natural brain dynamics. Front Hum Neurosci 8:444, (2014).

35. Luck SJ, Kappenman ES Oxford Handbook of Event-Related Potential Components (Oxford University Press, USA, New York) Available at: http://replace-me/ebraryid=11304148 (2011).

36. Kornhuber HH, Deecke L Brain potential changes in voluntary and passive movements in humans: readiness potential and reafferent potentials. Pflügers Arch - Eur J Physiol 468(7):1115–1124, (2016).

37. Brunia CHM CNVand SPN: Indices of Anticipatory Behavior. The Bereitschaftspotential (Springer US, Boston, MA), pp 207–227, (2003).

38. Di Russo F, et al. Beyond the “Bereitschaftspotential”: Action preparation behind cognitive functions. Neurosci Biobehav Rev 78:57–81, (2017).

39. Bozzacchi C, Giusti MA, Pitzalis S, Spinelli D, Di Russo F Awareness affects motor planning for goal-oriented actions. Biol Psychol 89(2):503–514, (2012).

40. Diener C, Kuehner C, Brusniak W, Struve M, Flor H Effects of stressor controllability on psychophysiological, cognitive and behavioural responses in patients with major depression and dysthymia. Psychol Med 39(1):77–86, (2009).

41. Elbert T, Rockstroh B, Lutzenberger W, Birbaumer N Slow brain potentials after withdrawal of control. Arch Psychiatr Nervenkr 232(3):201–214, (1982).

42. Casement MD, et al. Anticipation of affect in dysthymia: behavioral and neurophysiological indicators. Biol Psychol 77(2):197–204, (2008).

43. Bradley MM, Lang PJ Measuring emotion: the Self-Assessment Manikin and the Semantic Differential. J Behav Ther Exp Psychiatry 25(1):49–59, (1994).

44. Gramann K, Ferris DP, Gwin J, Makeig S Imaging natural cognition in action. Int J Psychophysiol 91(1):22–9, (2014).

45. Kothe C LabStreamingLayer. Available at: https://github.com/sccn/labstreaminglayer (2014).

46. Delorme A, Makeig S EEGLAB: an open source toolbox for analysis of single-trial EEG dynamics including independent component analysis. J Neurosci Methods 134(1):9–21, (2004).

47. Palmer JA, Kreutz-Delgado K, Makeig S AMICA: An Adaptive Mixture of Independent Component Analyzers with Shared Components Available at: https://sccn.ucsd.edu/~jason/amica_a.pdf (2011).

48. Bozzacchi C, Spinelli D, Pitzalis S, Giusti MA, Di Russo F I know what I will see: action-specific motor preparation activity in a passive observation task. Soc Cogn Affect Neurosci 10(6):783–789, (2015).

49. van Boxtel GJM, Böcker KBE Cortical Measures of Anticipation. J Psychophysiol 18(2/3):61–76, (2004).

50. Klein C, Rockstroh B, Cohen R, Berg P Contingent negative variation (CNV) and determinants of the post-imperative negative variation (PINV) in schizophrenic patients and healthy controls. Schizophr Res 21(2):97–110, (1996).

51. Barrett LF How emotions are made: the secret life of the brain (Houghton Mifflin Harcourt, New York) (2017).

52. Barrett LF, Bar M See it with feeling: affective predictions during object perception. Philos Trans R Soc B Biol Sci 364(1521):1325–1334, (2009).

53. Hillyard SA, Anllo-Vento L Event-related brain potentials in the study of visual selective attention. Proc Natl Acad Sci U S A 95(3):781–7, (1998).

54. Posner MI, Dehaene S Attentional networks. Trends Neurosci 17(2):75–9, (1994).

55. Mangun GR, Hopfinger JB, Heinze H-J Integrating electrophysiology and neuroimaging in the study of human cognition. Behav Res Methods, Instruments, Comput 30(1):118–130, (1998).

56. Gramann K, Töllner T, Müller HJ Dimension-based attention modulates early visual processing. Psychophysiology 47(5):968–78, (2010).

57. Gramann K, Toellner T, Krummenacher J, Eimer M, Müller HJ Brain electrical correlates of dimensional weighting: An ERP study. Psychophysiology 44(2):277–292, (2007).

58. Friston KJ, et al. Dopamine, Affordance and Active Inference. PLoS Comput Biol 8(1):e1002327, (2012).

59. Friston KJ Active inference and free energy. Behav Brain Sci 36(3):212–213, (2013).

60. Zumthor P Atmospheres (Birkhäuser, Basel) (2006).

61. Rockstroh B, Cohen R, Berg P, Klein C The postimperative negative variation following ambiguous matching of auditory stimuli. Int J Psychophysiol 25(2):155–167, (1997).

62. Gauthier P, Gottesmann C Communications in Electroencephalography. Electroencephalography and Clinical Neurophysiology (Elsevier), pp 534–535, (1977).

63. Kathmann N, Jonitz L, Engel RR Cognitive Determinants of the Postimperative Negative Variation. Psychophysiology 27(3):256–263, (1990).

64. Klepeis N, et al. The National Human Activity Pattern Survey (NHAPS): a resource for assessing exposure to environmental pollutants. J Expo Anal Environ Epidemiol 11(3):231–252, (2001).

65. Picard N, Strick PL Activation of the Supplementary Motor Area (SMA) during Performance of Visually Guided Movements. Cereb Cortex 13(9):977–986, (2003).

66. Adams RA, Shipp S, Friston KJ Predictions not commands: active inference in the motor system. Brain Struct Funct 218(3):611–643, (2013).

67. Zeki S Art and the Brain. Dædalus 127(2):71–103, (1998).

